# A completely resolved phylogenetic tree of British spiders

**DOI:** 10.1101/2021.03.12.434792

**Authors:** Rainer Breitling

**Affiliations:** Faculty of Science and Engineering, University of Manchester, M1 7DN Manchester, UK

## Abstract

The recent accumulation of increasingly densely sampled phylogenetic analyses of spiders has greatly advanced our understanding of evolutionary relationships within this group. Here, this diverse literature is reviewed and combined with earlier morphological analyses in an attempt to reconstruct the first fully resolved phylogeny for the spider fauna of the British Isles. The resulting tree highlights parts of the group where data are still too limited for a confident assessment of relationships, proposes a number of deviations from previously suggested phylogenetic hypotheses, and can serve as a framework for evolutionary and ecological interpretations of the biology of British spiders, as well as a starting point for future studies on a larger geographical scale.

## Introduction

In recent years, the number of large-scale phylogenetic analyses of spiders has been rapidly increasing. Molecular studies, as well as classical morphological work and integrative “whole-evidence” analyses, are covering large parts of spider diversity with increasing density. The last years have seen the publication of several comprehensive phylogenetic studies of the entire order, based on continuously increasing species coverage and ever-larger amounts of (mostly molecular) data (e.g., Agnarsson et al. 2013; Bond et al. 2014; Dimitrov et al. 2017; Fernandez et al. 2014, 2018; Garrison et al. 2016; Hedin et al. 2019; Kulkarni et al. 2020; Opatova et al. 2020; Ramírez 2014; Ramírez et al. 2019, 2021; Shao & Li 2018; Wheeler et al. 2017). Subsets of the order, from superfamilies to individual groups of genera, have also been the target of various analyses (e.g., Crews et al. 2020; Godwin et al. 2018; Hedin et al. 2018; Kallal et al. 2020; and numerous publications cited below for individual families). In addition, “DNA barcode” projects have provided a plethora of molecular genetic data for a wide range of spider species (e.g., Astrin et al. 2016; Blagoev et al. 2016), which can serve to complement earlier morphological analyses in an attempt to resolve phylogenetic relationships, especially within spider genera (Breitling 2019b,d).

It would be interesting to see how all these studies in combination can inform our understanding of the evolutionary relationships within a local spider fauna. Most ecological and faunistic work on spiders is done at a local level and would benefit from a clear and explicit evolutionary framework, i.e., a phylogenetic tree of the local spider fauna. While such a tree would be pioneering for arachnology, megatrees for local floras and faunas have been successfully constructed for many other groups, from European tetrapods (Roquet et al. 2014) and butterflies (Wiemers et al. 2020), to the vascular plants of the British Isles, Germany, the Netherlands, and Switzerland (Durka & Michalski 2012), as well as for British birds (Thomas 2008). They are considered an essential ingredient for evolutionary informed studies in ecology and conservation science (Roquet et al. 2013), e.g., providing the necessary phylogenetic framework for understanding the evolution of egg shell pigmentation in British birds (Brulez et al. 2016), or for identifying the effect of species’ traits on their population changes (Sullivan et al. 2015).

The general benefits of a phylogenetic framework are obvious: just imagine what ecological or faunistic studies would look like if all phylogenetic information were discarded: families and genera could not longer be used to structure the information. The ubiquitous pie charts that present survey results according to spider family would disappear. It would no longer be possible to state that the relative abundance and diversity of linyphiids increases towards higher latitudes, or that lycosids dominate in a pitfall sample. The resulting challenges would clearly be wide-ranging.

A fully resolved phylogenetic tree provides the same kind of information as the family and genus assignments, but with much finer granularity and without being restricted to arbitrary taxonomic categories. Ecological and faunistic studies can ask the same kind of questions about each of the clades in the tree that they would routinely ask about families or genera. This is a rich opportunity – and, importantly, the usefulness of a tree for this kind of analysis would not increase if it included a *global* set of species, but it would decrease if any *local* representatives were missing or if it were less than fully resolved.

Moreover, applying the phylogenetic results at a local level should help identifying gaps in our current understanding of the spider Tree of Life: is it even possible to plausibly reconstruct all the evolutionary relationships within a local spider fauna given the currently available data? Here, I attempt to answer this question for one particularly well-studied spider fauna, that of the British Isles. Great Britain and Ireland have a long and distinguished history of arachnological research. The fact that much of their fauna and flora was acquired after the glaciations of the ice ages has resulted in a relatively impoverished, but still interestingly diverse, spider fauna, which largely consists of species that are common and widespread across the Palaearctic. As a result, the vast majority of British spiders has been studied, illustrated and described repeatedly in great detail, providing an excellent starting point for determining their relationships. Many of the British species or their close relatives have been included in published phylogenetic studies, and a considerable fraction has been the target of comprehensive DNA barcoding projects. The systematics of the British spiders is reasonably stable, and their nomenclature and classification provide another, implicit source of information about the phylogenetic relationships within the group.

But isn’t it somewhat pointless to reconstruct a phylogenetic tree for such a small, stochastic subset of global biodiversity? Certainly not. For a start, all published phylogenetic hypotheses refer to limited subsets of global spider richness; a complete fully resolved tree of all 40,000+ spider species is a distant dream – and if it were constructed today, it would still lack all the undiscovered and most of the extinct species that are equally valid members of the great Tree of Life. Compared to the typical trees shown in the arachnological literature, the tree presented here is exceptionally large and densely sampled. But, more importantly, for the reasons listed above, the British spider fauna is far from a random subset of spider diversity: as a result of ecological history, it is strongly enriched in abundant, generalist and widespread species (and this still holds true for its recent arrivals). And, as a result of arachnological history, it is also strongly enriched in well-studied, carefully described and comprehensively analysed species, all of them repeatedly revised and illustrated and many of them barcode-sequenced; it is furthermore strongly enriched in type species of globally distributed genera, and type genera of many globally important families. Consequently, the resulting phylogenetic tree contains just the kind of species that will make it useful as a point of reference for expanding the tree, first to the Palaearctic fauna, and in the future to phylogenies on a global scale (even if these will more likely be generated from scratch – the synthesis of the literature provided here should give a first idea of the work still required to achieve this ultimate ambition).

Yet, a concern could remain that the geographically defined scope of the tree is inappropriate: after all, phylogeny does not respect geography, spiders do not know national borders, and the British spider fauna has certainly not evolved *in situ*, but is merely a part of global spider biodiversity, closely related to that of the nearby continent. However, this is also true for the many other taxa where local phylogenetic trees have proven to be of tremendous value. I would, moreover, argue that nobody doubts the value of checklists of the local spider fauna; for British spiders, such checklists are published regularly, with geographic scales ranging from individual nature reserves, to single counties and the entire country. Knowing which species are present in a local area provides the necessary context for any observations on individual groups on the list. It would be considered bizarre if such a list were sorted alphabetically by species epithet: at the bare minimum, the species will be grouped according to genus and family. In the case of British spiders, there has in fact been a longstanding tradition to arrange checklists even further by phylogenetic affinities, reflecting presumed evolutionary relationships in the order of species within genera, genera within families, and families within Araneae in general. A fully resolved phylogenetic tree takes this idea only a small and natural step further, making the proposed relationships explicit, unambiguous and testable.

As a purely intellectual exercise a tree like this meets an elementary scientific desire for taxonomic order based on evolutionary relationships and with the maximum possible resolution. But it also fulfils an important role as a didactic and mnemonic aid: it is much easier to learn and remember the characters and features of the members of a local spider fauna, if one is aware of their precise evolutionary relationships. In contrast to a determination key or other pragmatic arrangements of the species, a phylogenetically-informed “mind map” (i.e., a phylogenetic tree) is ready to grow by adding new species whenever needed, for instance because the taxonomic or geographic scope is expanded.

Of course, not every arachnologist will be interested in the detailed evolutionary relationships of their study animals; some may be satisfied with the coarse-grained picture provided by the Linnaean taxonomic hierarchy. But, the majority of curious naturalists will want to know how the species in their local patch are related to one another; initially, this interest will be quite independent of any relationships to species in other parts of the world. The tree identifies specifically the local sister group of each species or group of species, without being restricted by the arbitrary levels of the Linnaean hierarchy. It allows examining whether ecological, morphological or behavioural traits studied in a British spider community correlate with their patterns of evolutionary relatedness. But its usefulness doesn’t end there; even if this local tree only represents a subset of the global spider biodiversity, it would be applicable to studies across much of Northwest Europe, with little or no modification, and could easily be expanded to the spider fauna of large parts of Central Europe.

The central data source of this paper is a comprehensive synthesis of the published literature. As a result, the amount of original data will be limited, and a conscious effort was made to minimise the reliance on novel unpublished pieces of evidence. This turned out to be far from trivial: the emerging phylogenetic literature is surprisingly challenging to apply on a local level. Even the largest studies will include only a tiny fraction of the entire spider diversity; individual studies only partly overlap in the species included; and while there is a trend towards consolidation in some areas of the phylogenetic trees, there remain numerous parts where different studies arrive at widely different and inconsistent results, which are difficult to reconcile. Below, these challenges are discussed for selected individual examples, emphasising the necessarily subjective nature of some of the decisions made, in view of the still incomplete evidence available.

## Material and Methods

The phylogenetic reconstruction presented here is based on a synthesis of a wide range of published phylogenetic analyses, supplemented by the data in the taxonomic literature on British spiders, as well as additional barcode data from public databases. Given the heterogeneity of the available information, no rigid, formal approach could be expected to resolve the remaining conflicts and uncertainties convincingly. The construction of the preferred tree consistent with the cited data was done manually, and, in general, the results of studies including a denser sample of species in the relevant part of the tree were preferred over those with a sparser coverage; multi-gene molecular studies were preferred over purely morphological analyses; integrated studies, combining molecular and morphological data, over those using only one data type; studies with a larger number and diversity of molecular markers were preferred over those studying fewer genes; and barcode data from the literature and the BOLD database (Ratnasingam & Hebert 2007) were only used when they were not directly contradicted by the morphological evidence. In addition to recently published phylogenetic analyses, the full range of phylogenetic hypotheses implicitly or explicitly proposed in the traditional taxonomic literature was considered, as were all published morphological data, although no formal cladistic analysis of such data was attempted.

Morphological data for linyphiid spiders were obtained from Anna Stäubli’s identification key at the Spiders of Europe website (Stäubli 2020; https://araneae.nmbe.ch/matrixlinkey). The character states for *Nothophantes horridus* were added on the basis of Merrett & Stevens (1995, 1999), and those for the male of *Pseudomaro aenigmaticus* on the basis of unpublished observations by A. Grabolle. Data on the distribution of British spiders at the hectad level were downloaded from the website of the Spider and Harvestman Recording Scheme website (http://srs.britishspiders.org.uk/portal.php) and mapped onto the phylogenetic tree. Subsequently, for each of the subtrees, a Wilcoxon rank sum test was applied to test for significant differences in the value of ecological variables of interest for the species within the subtree (clade), compared to the rest of the species. Uncorrected p-values are reported, but these are all significant at the 0.05 level after Bonferroni correction for multiple testing based on the number of subtrees. Trees were visualised using the iTOL web tool (https://itol.embl.de/; Letunic & Bork 2019). Data were processed using custom-made R scripts, which are available from the author upon request.

In this article, the focus is on proposing a single fully resolved tree, expressing a single testable phylogenetic hypothesis for each triplet of species in the tree. Obviously, as will be mentioned repeatedly in the following discussion, not all of these hypotheses will be proposed with equal confidence. In many cases, alternative proposals would seem equally plausible. By unequivocally identifying one preferred hypothesis in each case, the proposed tree might be considered overly audacious. The advantage of this approach is that it facilitates future discussion and provides an unambiguous point of reference for necessary improvements and corrections.

## Nomenclature

The list of British spiders was obtained from the Spiders of Europe website (araneae.nmbe.ch; Nentwig et al. 2020) in May 2020. The list uses the World Spider Catalogue (WSC) nomenclature, which should be referred to for the currently accepted names of the species involved, as well as the authorities and additional taxonomic references for each of them. However, in the tree itself, the nomenclature has been adjusted to ensure that all genera are monophyletic, according to the proposed phylogenetic hypotheses. This renaming, which applies to a small minority of cases, largely follows the results of Breitling (2019a,b,c,d,e), with a few additional new combinations based on more recent published analyses.

### Subgenera

In a few cases, subgenera are explicitly proposed for traditional monophyletic groups within larger genera. Subgenera are currently rarely used in arachnology. This was not always the case and seems to be largely a historical consequence of the way the most widely used spider catalogues are organised; e.g., the WSC almost always ignores subgenus divisions, leading to justified concerns that re-classification proposals below the genus level might easily be overlooked. However, the use of subgenera increases the information content of a classification considerably (Wallach et al. 2009), and it avoids the instability caused by subdividing homogeneous and obviously monophyletic groups into smaller and smaller genera. Many of the most acrimonious taxonomic debates of recent years could probably have been avoided if subgenera had been more widely adopted as a useful category in spider systematics.

### Semispecies

Barcoding data have regularly revealed that closely related spider species share mitochondrial DNA haplotypes (Astrin et al. 2016; Blagoev et al. 2016; Domènech et al. 2020; Ivanov et al. 2018; Lasut et al. 2015; Nadolny et al. 2016; Oxford & Bolzern 2018). This has often been dismissed as the result of a “barcode failure”, potentially due to incomplete lineage sorting of recently diverged species. However, this explanation is clearly untenable for the majority of cases in spiders: almost always, the sibling species involved are highly mobile and very common ecological generalists that are sympatric and regularly syntopic over huge areas, often entire continents. Assuming a traditional allopatric model of speciation (Kraus 2000), this argues strongly against a recent divergence between the species.

Alternatively, it has been suggested that the lack of an interspecific barcode gap is the result of mitochondrial introgression, potentially facilitated by *Wolbachia*-mediated gene drives. This may well be the case; however, for such a scenario to be as common as it appears in spiders requires regular fertile hybridization between the species involved, with minimal negative fitness costs for the hybrid individuals. It has been suggested that such introgression would have negligible effects on the stability of the species boundaries, as the nuclear genetic contribution of one of the parents would be rapidly lost, only the maternally inherited mitochondrial genome remaining as a trace of the hybridisation event (Oxford 2019). This is, however, not correct: in fact, assuming a stable population size, it is the mitochondrial contribution that will be rapidly lost stochastically, but due to recombination events, at least some of the alleles from both parents will remain present in the nuclear genome of backcrossing generations for a long time. Some studies that have examined examples of a missing barcode gap in more detail have indeed found that at least some nuclear markers equally fail to differentiate the species involved (Lasut et al. 2015; Spasojevic et al. 2016). The exchange of nuclear alleles is not necessarily obvious in the form of a gradient of phenotypically intermediate individuals, if backcrossed hybrids rapidly approach the phenotype of one of the parents (Oxford 2019).

The relevance of this phenomenon for the present analysis is that species that exchange mitochondrial DNA barcodes are likely to also exchange favourable (or neutral) nuclear alleles regularly (examples are known from crop pests exchanging pesticide resistance genes, but also from mimicry complexes in butterflies; e.g., Valencia-Montoya et al. 2020 and Zhang et al. 2016). Such groups do not form independent evolutionary individuals yet – and they are not mutually monophyletic –, and consequently it would be meaningless to propose a fully resolved phylogenetic tree for them. This is only of minor import for species pairs, but in several of these cases, especially among wolf spiders, groups of three or more species are involved globally.

In the tree presented here, I propose to apply the semispecies concept to such cases. Semispecies are groups of organisms in the “grey zone” of speciation, which have completed some of the necessary steps towards full species separation, but not all of them. Semispecies have often been used to classify island populations (as a synonym of allospecies), and the rare examples in arachnology also applied the concept strictly to allopatric populations (Kraus 2000); however, such a narrow interpretation is not necessary, and in groups with a more mature taxonomy (especially birds and butterflies) locally sympatric (as well as parapatric) groups of semispecies are identified with some regularity (e.g., Helbig et al. 2002; Smith et al. 2010), albeit not without lively debate regarding individual cases (e.g., Pfander 2011; Vane-Wright 2020).

Importantly, semispecies are not by definition species *in statu nascendi*; their evolutionary future cannot be predicted with certainty. In some cases, it might be that disruptive selection, reinforcement and related mechanisms will allow them to progress to full speciation in sympatry. But it is equally possible that they will continue to maintain gene flow indefinitely or even merge into a single homogeneous freely panmictic population. Interestingly, among members of the *Eratigena atrica* species group, individuals in the overlapping part of the range show a tendency towards intermediate phenotypes (Oxford 2019), in contrast to the expectation of “character displacement”, which would predict that sympatric populations would become more clearly distinct phenotypically, e.g., to minimise harmful hybridisation or ecological niche overlap.

In the phylogenetic tree, semispecies are explicitly identified by including a “superspecies” name, i.e., the name of the first member of the group to be described, in brackets before the species epithet. This highlights areas of the tree where speciation may not yet be complete, and where the question “one species or two (or three)?” is ill posed. Studies such as the work of Ivanov et al. (2018), Domènech et al. (2020), and Oxford (2019) show that each of these cases will be a rich ground for future evolutionary insights.

## Results

The deep phylogenetic relationships of spider families have recently been addressed by a series of molecular genetic studies (Agnarsson et al. 2013; Bond et al. 2014; Dimitrov et al. 2017; Fernandez et al. 2014, 2018; Garrison et al. 2016; Opatova et al. 2020; Ramírez 2014; Ramírez et al. 2019, 2021; Shao & Li 2018; Wheeler et al. 2017). These show considerable agreement with previous morphological analyses (e.g., the trees shown in Coddington 2005 or Jocqué & Dippenaar-Schoeman 2006), and where they disagree (e.g., regarding the non-monophyly of orb-web weavers or the placement of Mimetidae among Araneoidea), the strong statistical support for the alternative hypotheses and the agreement of independent molecular analyses indicate that the latter recover the true evolutionary relationships. Overall, the last few years have seen rapid convergence towards a stable consensus, although a few details, such as the sister group of Salticidae, remain somewhat contentious. In addition to the British species, the tree in **Figure 1** includes all families known to be present in Europe, to provide a broader context for the phylogenetic position of the British fauna.

**Figure 1.**
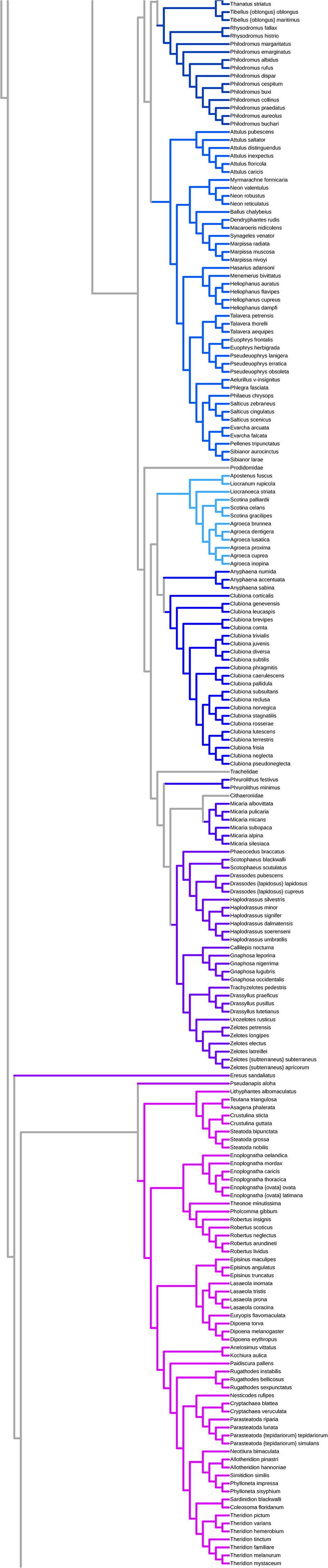
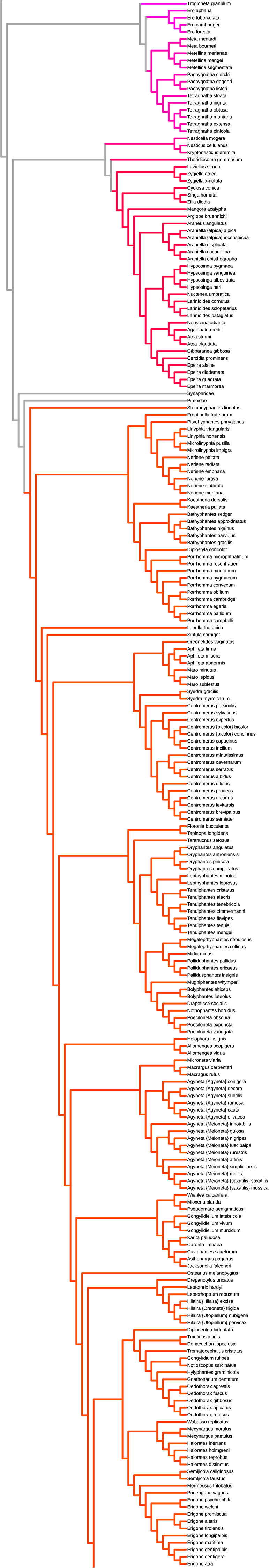
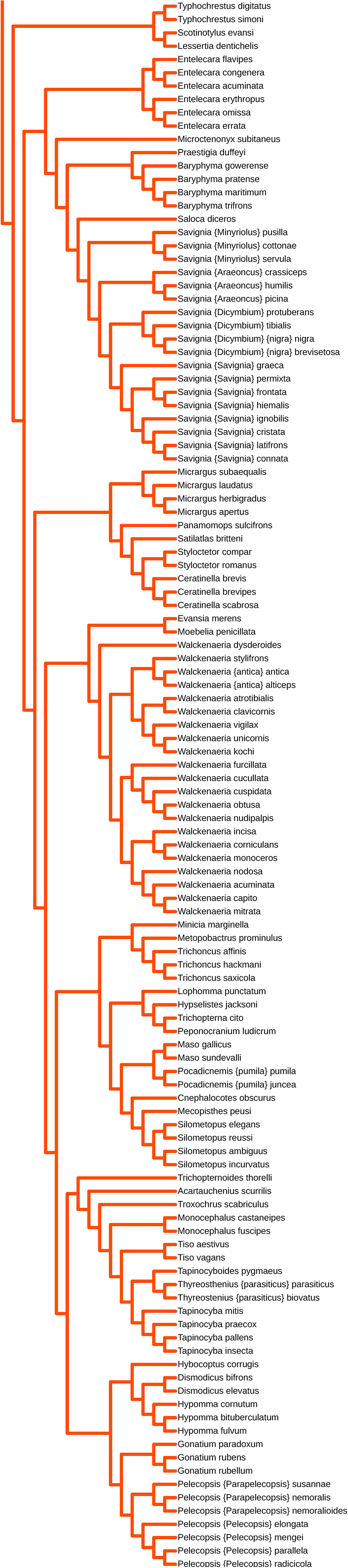
Fully resolved phylogenetic tree of the British spider species. All non-British spider families with European representatives are included for reference, and the tree is rooted using Liphistiidae as the outgroup. Branch lengths are arbitrary and do not indicate the timing or degree of divergence. For readability, species from the same family are shown in the same colour.

While establishing the family-level framework is by now relatively easy, reconstructing the phylogenetic relationships within each family required the reconciliation of a diverse range of published proposals and personal judgement calls, on a case-by-case basis. Once a monophyletic group of *N* species has been established, *N* – 2 decisions (justifying *N* – 2 branch points) are required to reconstruct a fully resolved phylogenetic tree. For the more than 600 British species in the present tree, obviously, not all of these decisions can be elaborated in detail here. Importantly, the evidence supporting each decision (including published cladograms, character matrices, illustrations of the genitalia, and a variety of DNA sequences) could not be presented in the text; doing so would require reproducing a large fraction of the taxonomic literature on British and European spiders. Instead, the reader is referred to the data in the cited literature. However, selected examples from each family with more than 2 members are discussed in alphabetical order below.

### Agelenidae

The tree for this family is based on the detailed study by Bolzern et al. (2013), combining morphological and molecular data for a dense sample of species. The deep relationships of the genera show major differences compared to the analysis of Wheeler et al. (2017), which included individual representatives of the same genera. Crews et al. (2020) include a larger number of species, and recover a tree that is similar to that reported by Wheeler et al. (2017), but seems overall poorly resolved. These disagreements illustrate how fragile some of the results of even the most recent molecular analyses still can be. The arrangement proposed by Bolzern et al. (2013) is preferred, as the molecular results in this case agree quite closely with those of a morphological analysis, while in the trees presented by Wheeler et al. (2017) and Crews et al. (2020) in particular the placement of *Coelotes* deeply within the remaining Agelenidae *s. str*. is unexpected (the genus is typically placed in the subfamily Coelotinae, which is sometimes even considered a separate family Coelotidae). *Agelena longipes* is a phantom species as defined by Breitling et al. (2015, 2016), i.e., it was not rediscovered since its original description in 1900. It is thus considered a *nomen dubium* and is not included in the tree.

### Amaurobiidae

The topology of the tree within this small group is determined by the morphological affinities, and confirmed by available barcode data for all three species.

### Anyphaenidae

The relations in this case are inferred on the basis of the similarities in pedipalp and epigynum of *Anyphaena accentuata* and *A. sabina*, as well as the rather close similarity the barcodes for these two species, which indicate that they are probably are pair of sister species, while *A. numida* is more distantly related.

### Araneidae (and Zygiellidae *sensu* Wunderlich 2004 = Phonognathidae *sensu* Kuntner et al. 2019)

The relationships of species in this group are based on the analyses of Kallal & Hormiga (2019), Kallal et al. (2020), and Scharff et al. (2020). The genus-level backbone closely follows the results of Scharff et al. (2020), as this study has the densest coverage of relevant species. Kallal et al. (2020) differ only in details. The internal topology within *Araniella* follows Spasojevic et al. (2016); for other genera the relationships are informed by barcode data, which are available for all species. The placement of *Zilla* is only very weakly supported; it is based on Tanikawa’s assessment that this genus is closely related to *Plebs/Eriophora* (Tanikawa 2000), which in the tree of Scharff et al. are closely related to *Singa*.

The genus *Araneus* in the traditional sense is clearly polyphyletic. The placement of *Araneus* (*s.str*.) *angulatus* is based on barcode similarity to *A. bicentenarius*, which is placed apart from *A. diadematus* and its relatives in Scharff et al. 2020. The close relationship between *A. angulatus* and *A. bicentenarius* is confirmed by their morphological similarity, which led earlier authors to consider *A. bicentenarius* a variant of *A. angulatus* (Levi 1971). The available genus name for the *A. diadematus* group would be *Epeira*, and this is used in the tree.

The placement of the *Atea* species is unclear. Their barcode sequences indicate that they are not closely related to *A. angulatus* or the *A. diadematus* group of the highly polyphyletic *Araneus* s. *lat*.; as the genus *Atea* has historically often been placed close to *Agalenatea*, the two British species are placed there, but with some reluctance, as there seems to be no convincing evidence for a close relationship between the two genera.

### Cheiracanthiidae

Relationships within this small group remain obscure. Wolf (1991) discusses *Cheiracanthium pennyi* and *C. erraticum* as being more similar to each other than *C. erraticum* and *C. virescens*, contra Helsdingen (1979). Barcodes for *C. pennyi* are not available, so the molecular data cannot help resolving the case. The pedipalp without ridge and the epigynum with short insemination ducts set *C. pennyi* apart, but these could be autapomorphic. The striking red dorsal line of the opisthosoma is shared by *C. pennyi* and *C. erraticum*, but here the loss in *C. virescens* could be autapomorphic. The preferred topology represented in the tree gives precedence to the genital similarities, but without a sound cladistic basis for this decision.

### Clubionidae

The backbone of the topology within this family largely follows Breitling (2019c). *Microclu-biona sensu* Wunderlich (2011), i.e., the *trivialis* group *sensu* Mikhailov (1995) is considered monophyletic (as in some of the barcode analysis results). *Euryclubiona sensu* Wunderlich (2011) is also considered monophyletic, and is recovered as such in most of the barcode results; the internal topology within this group is based on morphological affinities. For the reasons discussed in Breitling (2019d), none of these subgenera is formally recognised here, but a future subdivision of *Clubiona* into well-defined *sub*genera would certainly be highly preferable over a splitting of this homogeneous and obviously monophyletic group into separate genera.

The position of *C. rosserae* as sister of *C. stagnatilis* is based on the original description and Wiehle (1965), while the position of *C. subtilis* and *C. juvenis* is based on morphological similarities to their proposed sister species. *C. corticalis* is placed basally within the genus, in agreement with its unique morphology and most barcode analyses.

*C.facilis*, which is also listed as a member of the British spider fauna is possibly a synonym of *C. phragmitis*, based on a malformed specimen, analogous to the case of *Philodromus depriesteri* discussed in Breitling et al. (2015). The close morphological similarity to *C. phragmitis* discussed in the original description (sub *C. holosericea*), as well as the “atrophic” appearance of the epigynum in the accompanying illustration, would seem to support this interpretation. Examination of the type material in the Pickard-Cambridge collection in the Oxford University Museum of Natural History (Bottle 2312.1) could provide further insights, but for now the name is considered a *nomen dubium* and is not included in the tree.

### Dictynidae

Dictynidae and Hahniidae are closely related, and the position of some genera in either of them remains unclear. According to Crews et al. (2020), both *Cicurina* and *Lathys* “remain unplaced”. For now, and for the purposes of the preferred tree, placement of *Lathys* as basal to the remaining Dictynidae and *Cicurina* basal to Hahniidae seems most justified, as this is roughly compatible with the results presented by Wheeler et al. (2017) and consistent with at least one of the analyses shown in Crews et al. (2020). In the analysis by Crews et al. (2020), *Dictyna* was recovered as polyphyletic, but here it is assumed that the British representatives form a monophyletic group, as do the other genera in these two families.

The placement of *Altella* is questionable. Wunderlich (1995a, 2004a) considers the genus a junior synonym of *Argenna*, indicating at least a close relationship between the two genera. But in the same works he also considers *Brigittea* (and *Emblyna*) as synonyms of *Dictyna*; for this to be correct, it would be necessary to considerably expand the scope of *Dictyna*, to also include the quite distinct genus *Nigma*.

Given the close morphological similarities (and barcode similarities) between *Argenna*, *Altella* and *Dictyna* s.lat., *Argyroneta* is placed basal to a clade uniting these two groups, despite results by Crews et al. (2020) that argue for joining *Argenna* and *Argyroneta* instead.

The internal topology within *Nigma* is supported by morphological affinity that indicates a sister group relationship between *N. puella* and *N. flavescens*. The same argument applies, with less conviction, in the case of *Lathys*. The topology within *Dictyna s. str*. is based on a combination of morphological similarities and barcode data for a subset of the species.

### Dysderidae

Platania et al. (2020) show that *Harpactea* in its present form is a polyphyletic assembly of unrelated groups, the two British species being placed in deeply separated clades. Non-monophyly is also found for several other genera of Harpacteinae (*Folkia*, *Dasumia*), as well as for some of the previously proposed species groups within *Harpactea*. This indicates that the morphological recognition of monophyletic units within this family is unusually challenging. Consequently, instead of separating the two British species into different genera, it appears more pragmatic to extend the *Harpactea* genus concept and treat them as members of a single *Harpactea s. lat*.

### Gnaphosidae (and Micariidae *sensu* Mikhailov & Fet 1986)

The results of Wheeler et al. (2017) show that morphological data have so far failed to converge on a stable and reliable phylogenetic reconstruction for Gnaphosoidea. Recent morphological analyses by Rodrigues & Rheims (2020) and Azevedo et al. (2018) show fundamental differences compared to the molecular analysis presented by Wheeler et al. (2017). For example, they place Prodidomidae deep within Gnaphosidae; a placement that molecular analyses contradict with strong support. On the other hand, in the molecular analysis, traditional Gnaphosidae are highly polyphyletic.

At this point, a conservative tree will largely follow the morphological results and traditional arrangements. The morphological analyses do not fully resolve the relationship between the subfamilies Gnaphosinae, Zelotinae, and Drassodinae. The preferred arrangement at this level is based on the molecular data.

The placement of *Urozelotes* in the tree is tentative; it could with similar justification be placed as sister to *Drassyllus*+*Trachyzelotes*, rather than *Zelotes*. The internal structure of *Drassodes, Drassyllus, Haplodrassus* and *Zelotes* is based on a combination of morphological similarities and barcode data. In the case of *Haplodrassus*, the placement of *H. minor* and the deeper branches are ambiguous. The relationships within *Gnaphosa* are based on morphological similarities only.

Micariidae are here treated as a separate family (**stat. rev.**), sister to Cithaeronidae, based on the results in Azevedo et al. (2018) and Rodrigues & Rheims (2020). This separation seems justified given the long-standing debate about the placement of *Micaria*, which often was included in Clubionidae instead of Gnaphosidae. Given the chaotic results for Gnaphosidae in Wheeler et al., this preference is obviously only weakly supported. The internal structure of the tree for the genus *Micaria* follows Breitling (2017). The placement of *M. albovittata* is based on Wunderlich’s inclusion of the species (sub *M. romana*) in the *pulicaria* group (Wunderlich 1980). The placement of *M. silesiaca* is based on its inclusion in the *silesiaca* group (Wunderlich 1980).

### Hahniidae

The inclusion of *Cicurina* in this family is discussed above under Dictynidae. The inclusion of *Mastigusa* is particularly weakly supported. The British species has formerly been placed in *Tetrilus*, *Tuberta*, and *Cryphoeca*. Its placement in Hahniidae is based on its presumed relationship to *Cicurina* (e.g., Murphy & Roberts 2015); Wunderlich (2004a) places it in Cryphoecinae instead. The two forms of British *Mastigusa* are here conservatively considered semispecies, rather than morphs of a single species, given their apparent ecological separation. However, if this interpretation is correct, the name of the *macrophthalmus* form would probably need to be changed, as the British specimens don’t seem to belong to the species originally described under this name from Eastern Europe (Wunderlich 1995b). The relationships between the genera are based on analysis of the barcode data available for the family, as is the internal topology of *Hahnia*. The internal topology of *Iberina* is based on morphological affinities, but as the male of *I. microphthalma* is unknown, this remains tentative.

### Linyphiidae

This family is particularly difficult to analyse, not just because it is the largest of the British spider families, but also because a large part of recent taxonomic work has been dedicated to splitting genera into poorly supported smaller units on the basis of typological arguments, instead of identifying convincing relationships between genera; together with the traditionally poor genus concepts in this group, this has created such a degree of confusion that even a considerable amount of detailed phylogenetic analyses (both molecular and morphological) have not been able to completely clarify the situation, and the phylogenetic relationships of many genera remain unresolved at all levels. Additionally, while the molecular and morphological analyses show some convergence in a few important areas of the tree, a large fraction of the published trees is still highly unstable, and the addition of new characters or species can lead to major rearrangements (see, e.g., the discussions in Miller & Hormiga 2004 and Paquin et al. 2008). The proposal advanced here can only be a very first attempt at providing a comprehensive phylogenetic hypothesis for this group (within strict geographical limits).

The framework for the proposed linyphiid phylogeny is provided by the molecular analyses of Wang et al. (2015) and Dimitrov et al. (2017). This is complemented by the increasingly comprehensive morphological analyses of the entire family or large subgroups in Duperré & Paquin (2007), Gavish et al. (2013), Hormiga (1993, 1994, 2000), Hormiga & Scharff (2005), Miller & Hormiga (2004), Paquin et al. (2008), and Sun et al. (2012). Most importantly, the relative placement of genera required a much larger degree of personal interpretation of the traditional taxonomic and morphological literature. The British Linyphiidae were comprehensively analysed in terms of pedipalp morphology by Merrett (1963) and Millidge (1977), and less comprehensively in terms of their female genitalia by Millidge (1984, 1993). This information was complemented by the phylogenetic assessments implicitly (and rarely explicitly) contained in the works of Wiehle (1956, 1960) and Roberts (1987), as well as a thorough assessment of the morphological data encoded in the interactive key of linyphiid species by Anna Stäubli (Stäubli 2020, http://www.araneae.nmbe.ch). The barcode analyses presented in Breitling (2019b) provided additional information, but were mostly used for determining the relationships within genera.

The internal topology of *Agyneta* is based on a careful interpretation of barcode data, in conjunction with a morphological analysis. *Agyneta* is a good example of a genus where species identification is challenging and the resulting mis-identifications cause difficulties in interpreting barcode database information. In the British fauna, *Meioneta* and *Agyneta* seem to be mutually monophyletic and could be maintained as subgenera, but in the global context, they should remain unified (together with a number of smaller genera) in *Agyneta s. lat*.

Following Breitling (2019b), *Saaristoa* is considered a junior synonym of *Aphileta*, and *Centromerita* a junior synonym of *Centromerus*. In both cases, the proposed phylogenetic hypotheses support this synonymy, as it is necessary to maintain the monophyly of all named genera.

*Collinsia* is treated as a junior synonym of *Halorates*, following Buckle et al. (2001), Millidge (1977), Roberts (1987), and Tanasevitch (2009). As the proposed tree shows, it would be impossible to maintain *C. inerrans* in the same genus as *C. holmgreni* / *C. distinctus*, if *H. reprobus* is excluded. Joining the two genera in *Halorates s. lat*. seems more conservative in the short run, than a splitting off of *C. inerrans* (in *Milleriana*), in the absence of a comprehensive revision of this and several related genera. The barcode data indicate a general confusion in this group, where most genera are not recovered as monophyletic. This is not fully reflected in the proposed tree, which gives priority to the morphological similarities; e.g., in its COI barcode, *Mecynargus paetulus* seems to be closer to *H. inerrans* than to the type of its genus, and *H. inerrans* closer to *M. paetulus* than to *H. holmgreni;* complementary information from a larger range of molecular markers would be required to justify such a major rearrangement.

*Dicymbium* is treated as a subgenus in a considerably expanded genus *Savignia*, resulting in a number of new combinations, as shown in the tree. This change in rank is consistent with earlier proposals by Millidge (1977) concerning the expansion of *Savignia* to include most of the members of his “*Savignya* genus group”. It is also supported by both molecular and morphological evidence as discussed in Frick et al. (2010) and Breitling (2019d). *Savignia* (*Dicymbium*) *brevisetosa* is certainly not a subspecies of *S*. (*D*.) *nigra* in the current sense, as the two occur sympatrically. The genitalia are indistinguishable and the two forms are not clearly ecologically distinct, although syntopic occurrence apparently is rare; it is therefore quite likely that they are synonymous, *brevisetosa* merely being a geographically restricted variant of the male prosomal morphology, as suggested by Roberts (1987). However, the genetic barcode data show two clusters (BINs) among the *Dicymbium* specimens, which could indicate the presence of two closely related species, one of which might correspond to the *brevisetosa* form, occasional intermediate specimens being the result of sporadic hybridisation. The two forms are therefore here considered conservatively as semispecies.

*Erigone maritima* is considered a separate species, distinct from *E. arctica s. str*., based on the considerable barcode gap between Nearctic and Palaearctic specimens identified as “*Erigone arctica” s. lat*. Whether the palaearctic species can be meaningfully subdivided into subspecies is currently an open question; given the high mobility and vast range of *Erigone* species, which are among the most frequent aeronauts, a relevant subspecific differentiation seems rather unlikely. Many of the morphologically well-defined *Erigone* species show a surprisingly narrow barcode gap, indicating relatively recent differentiation and arguing further against the probability of the existence of morphologically all but cryptic subspecies.

*Mermessus* (sub *Eperigone*) was considered as probably closely related and possibly the sister group of *Erigone s. lat*. by Millidge (1987), and the barcode data support this placement.

*Erigone longipalpis meridionalis* is a phantom species as defined by Breitling et al. (2015, 2016) and probably only represents intraspecific variation of *E. longipalpis*. It is thus considered a *nomen dubium* and not included in the tree.

*Frontinellina* is considered a junior synonym of *Frontinella*, because of the close genetic affinities between representatives of the two genera.

*Hilaira* is considered a senior synonym of *Oreoneta*. When separating *Oreoneta* from *Hilaira*, Saaristo & Marusik (2004) point out that *H. nubigena* and *H. pervicax* are also not conspecific with the type species of *Hilaira, H. excisa*. Instead of creating three poorly delimited genera, it is far more informative to consider the three groups as subgenera within a monophyletic genus *Hilaira s. lat*., sister to *Sciastes*. For the subgenus including *H. nubigena* and *H. pervicax*, the name *Utopiellum* (type species: *H. herniosa*) might be available, and it is here used in the tree.

*Maso* and *Pocadicnemis* are strongly united in the barcode data; their position relative to other higher erigonines is less clear. They are placed in the same group by Merrett (1963; Group E) and Locket & Millidge (1953; all tibiae with 1 dorsal spine; with Tm IV), but these are rather large groups, and the morphology of the two genera does not indicate a particularly close relationship to each other or other genera.

*Oryphantes* is considered a senior synonym of *Anguliphantes, Improphantes, Mansuphantes* and *Piniphantes*, following Breitling (2019b), and *Palliduphantes antroniensis* is also transferred to *Oryphantes s. lat*., where it belongs on the basis of its genital morphology (Bosmans in Heimer & Nentwig 1991), as confirmed by barcode information. As explained in Breitling (2019d), the synonymy is also supported by the observation of Wang et al. (2015) that a combination of a large number of genetic markers, including mitochondrial (COI and 16S) as well nuclear sequences (18S, 28S, H3), recovers *Anguliphantes* and *Oryphantes* as mutually polyphyletic with strong bootstrap support.

Millidge (1977) and Merrett (1963) point out similarities between *Ostearius* and *Dona-cochara*/*Tmeticus*, and Wiehle (1960) places *Ostearius* in his Donacochareae. However, this traditional placement of *Ostearius* in a clade with *Tmeticus* and *Donacochara* has long been dubious, and it is not supported by any of the recent analyses. Even the sister group relationship between the latter two is not strongly supported by any of the newer data. Hormiga (2000) and subsequent morphological assessments place *Tmeticus* far from *Ostearius*. The barcode data also do not indicate a close relationship: there, *Ostearius* is sister to *Eulaira*, matching Millidge’s earlier morphology-based proposal (Millidge 1984).

*Pelecopsis susannae* is transferred to *Parapelecopsis*, based on similarity of genitalia and absence of dorsal spines on its tibiae. As this indicates that the boundary between the two genera is not quite clear, they are here treated as subgenera of *Pelecopsis s.lat*., and in the global context *Parapelecopsis* should possibly be discarded altogether.

*Poeciloneta* is treated as a senior synonym of *Agnyphantes* and *Obscuriphantes*. While the necessity of this merger is not obvious in the context of the British fauna, where each of these genera is represented by a single species, the global analysis shows that this move is required to obtain a monophyletic genus *Poeciloneta*.

In the case of morphologically homogeneous genera, where even the species boundaries have long been ambiguous and species groups have been fluid at best, in the absence of genetic data the proposed phylogenetic relationships can be little more than a poorly educated guess. The genus *Porrhomma* is a good example of this situation. The preferred tree presented here is based on a rather subjective assessment of the morphological affinities of the included species.

The placement of *Pseudomaro* as sister of *Mioxena* is based on unpublished data on the morphology of the males (A. Grabolle https://wiki.arages.de/index.php?title=Pseudomaro_aenigmaticus) These indicate that the two genera may even be synonymous, but a formal synonymisation should await a formal publication of the description of male *Pseudomaro* specimens.

*Savignia* is here considered in the broadest sense, as discussed in Breitling (2019d). It includes the former genera *Dicymbium*, *Minyriolus*, *Glyphesis*, *Araeoncus*, *Diplocephalus* and *Erigonella*. Various earlier authors, including Bosmans (1996), Frick et al. (2010), Holm (in *lit*. in Millidge 1977), and Millidge (1977) had already found that this group is so homogeneous and the genera so poorly defined that they should probably be merged in a single genus. The barcode results confirm this assessment. The subgenus assignments try to identify monophyletic groups, at least within the context of the British fauna, but they are tentative only, given that no comprehensive global analysis of the genus group has been performed, and their practical value could be debated. *Savignia connata jacksoni* is considered an infrasubspecific variant of *Savignia connata*, following Roberts (1987), and is therefore not included separately in the tree.

**Figure 2** shows a mapping of selected morphological characters used for traditional “pragmatic” classifications of British linyphiids onto the proposed phylogenetic tree of this family.

**Figure 2.**
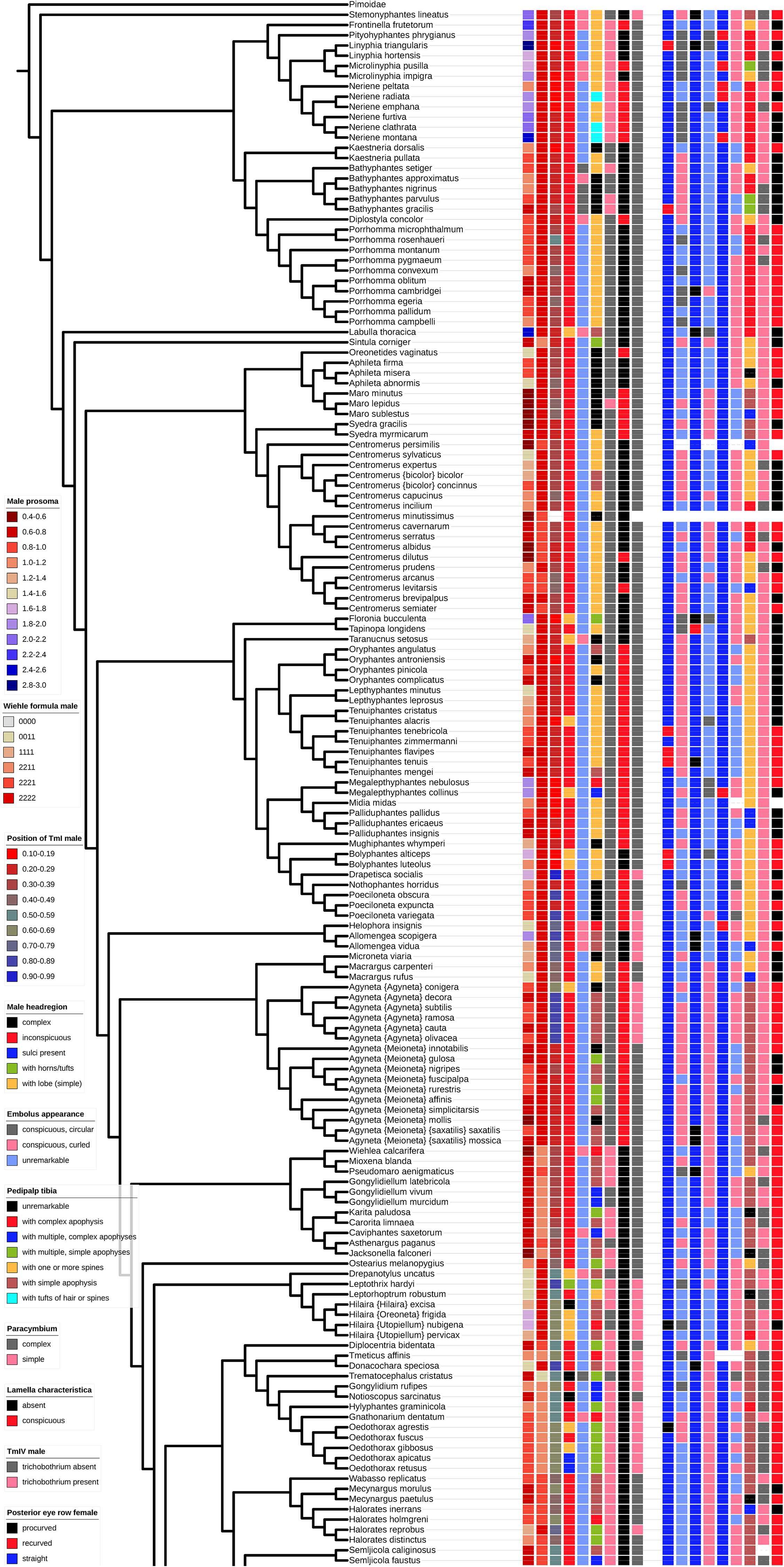
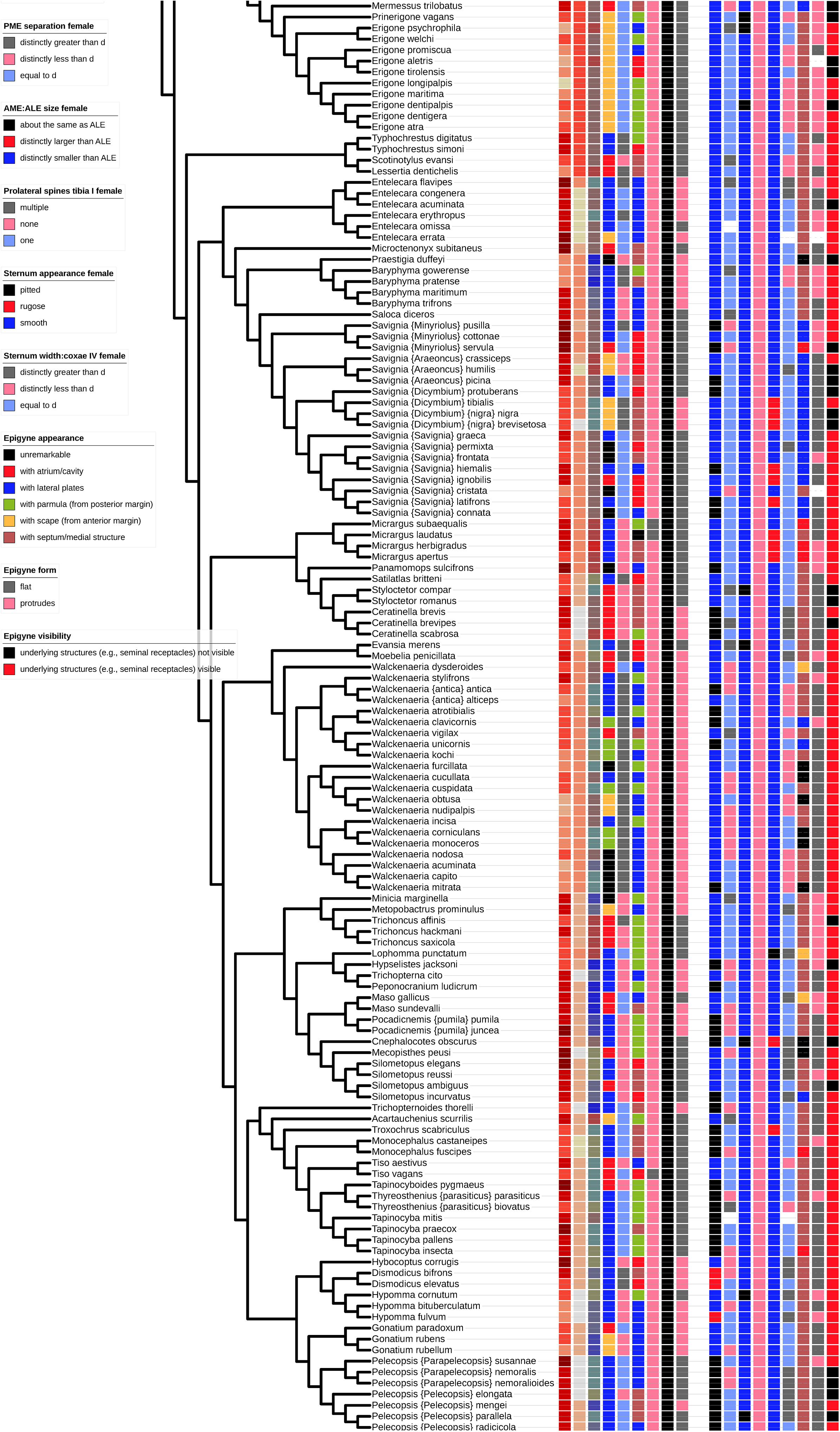
Selected morphological character states mapped onto the phylogenetic tree of British linyphiid spiders. The framework provided by the phylogenetic information allows the clear identification of patterns of character evolution, including a considerable degree of homoplasy for all characters. The definition of characters and character states is based on the linyphiid identification key by Anna Stäubli, provided on the Spiders of Europe website. The order of columns (from left to right) follows the order of the legends (from top to bottom). The left block of columns refers to characters applying to the male, the right to characters of the female. The female of *Centromerus minutissimus* has not yet been described.

### Liocranidae

The status and phylogeny of this family are controversial; morphological analyses by Bosselaers & Jocqué (2002) and Ramírez (2014) do recover the family as presently defined as strongly polyphyletic. The molecular results of Wheeler et al. (2017) agree. Only the morphological study of Bosselaers & Jocqué (2013), which analyses the densest sample of species, including all genera found in the British Isles, presents a monophyletic Liocranidae *s.lat*. The preferred tree presented here shows a compromise between the different analyses: while it proposes that the British representatives of Liocranidae are united in a monophyletic group, it modifies the arrangement of genera suggested by Bosselaers & Jocqué (2013) to match the observation by Ramírez (2014) that *Liocranum* and *Apostenus* are more closely related to each other than to *Agroeca* (which Ramírez wants to remove to Clubionidae). *Scotina* was not included in the study by Ramírez (2014), but is morphologically closer to *Agroeca*, although historically, the species of this genus have been placed in *Agroeca, Liocranum*, and *Apostenus* (*S. palliardii* in all three).

Relationships within *Scotina* are based on the phylogeny proposed by Bosselaers & Jocqué (2013). The arrangement within *Agroeca* follows Braun’s (1967) division of the genus into two species groups, (*A. lusatica*, *A. brunnea*, *A. dentigera*) vs. (*A. cuprea*, *A. proxima*). This contradicts the results of Bosselaers & Jocqué (2013), but is supported by barcode data. The placement of *A. lusatica* (sister to *A. dentigera*) and of *A. inopina* (sister to *A. cuprea*) is based on the stated morphological similarities in Grimm (1986).

### Lycosidae

The arrangement of genera in this family follows Piacentini & Ramírez (2019). To infer the relationships within *Trochosa*, the morphological results of Hepner & Milasowszky (2006) are complemented with barcode data for several of the species. Following Breitling (2019b), *Piratula* is considered a junior synonym of *Pirata*, although the genera traditionally included in this genus form a monophyletic group within the context of the British Isles, as supported by barcode data. The placement of *P. tenuitarsis* is based on its morphological similarity to *P. piraticus* (Kronestedt 1980).

The relationships within *Alopecosa* are based on morphological similarities and barcode results. The use of *A. barbipes*, instead of *A. accentuata*, for the British species is based on the argument presented by Breitling et al. (2016b), which was curiously misunderstood by Canard & Cruveillier (2019).

Arrangements within *Arctosa* follow the results of Knülle (1959), complemented by barcode data.

In *Pardosa*, the deep branches of the tree are inferred on the basis of morphological affinities, while barcode data resolve the internal relationships within species groups, including a number of semispecies complexes, which are particularly common in this genus.

### Mimetidae

The tree is based on the morphological assessments presented by Thaler et al. (2004), which are fully consistent with the barcode data. The placement of *Ero tuberculata*, which remains ambiguous in Thaler et al. (2004), is based on somatic and genital morphology, which suggests a closer relationship between *E. cambridgei* and *E. furcata*, than between either of them and *E. tuberculata*.

### Miturgidae

The backbone of the arrangement in this family is based on the barcode data available for three of the species, as the genus present in Britain is morphologically rather homogeneous. The placement of *Zora armillata* is based on its morphological similarity to *Z. spinimana*, implied in the determination keys presented by Urones (2005) and by Wunderlich in Heimer & Nentwig (1991). Of course, this is a rather weak argument, as these keys are intended as pragmatic aids to identification, not as statements of phylogenetic hypotheses.

### Nesticidae

Pavlek & Ribera (2017) illustrate the close relationship of *Nesticus* and *Kryptonesticus*, based on morphological and molecular data. *Nesticella* is morphologically quite distinct, and barcode data support the proposed arrangement.

### Oonopidae

Despite recent concerted efforts to expand our knowledge of oonopid biodiversity, most notably the Goblin Spider Planetary Biodiversity Inventory (http://research.amnh.org/oonopidae/), our understanding of the phylogenetic relationships within the group remains pitifully poor. Busschere et al. (2014) include the British genera in their analysis, but the resolution of their tree is low and the phylogeny preferred here has only low support. *Orchestina dubia* could be a considered a phantom species following the definition by Breitling et al. (2015, 2016). In contrast to other phantom species mentioned here, there remains a distinct possibility it will turn out to be a valid species, and it is therefore included in the tree for completeness.

### Philodromidae

The basic relationships between genera are based on Wheeler et al. (2017), while internal relationships are based on the data presented in Breitling (2019b).

The placement of *Philodromus buchari* is tentative, based on the morphological affinities indicated by Muster & Thaler (2004).

### Pholcidae

Compared to the similarly diverse Oonopidae, and despite a much smaller number of specialists working on the group, pholcid phylogeny has been studied extensively using molecular and morphological approaches and has reached reasonable stability. The relationships preferred here are based on Huber et al. (2018), but the placement of *Holocnemus pluchei* remains ambiguous.

### Salticidae

Salticidae are another family with a well-resolved phylogenetic backbone, based on morphological and molecular analyses (Maddison 2015; Maddison et al. 2014, 2017; Zhang & Maddison 2015). The deep relationships preferred here are mostly based on the results presented in Maddison (2015) and Maddison et al. (2017).

Relationships within *Euophrys* and closely related species are based on the results in Breitling (2019e). For *Sitticus* s.lat. the presented tree follows Maddison et al. (2020). The relationships within *Neon* are inferred on the basis of the morphological data in Lohmander (1945). The close relationship of *Synageles* and *Marpissa* is (very) weakly supported by their barcode sequences, as are the internal relationships within *Heliophanus*. The relationships of species within *Marpissa* and *Salticus*, in contrast, are robustly supported by the barcode data.

### Segestriidae

Morphologically (in terms of palp, endogyne, colour pattern, and leg spines), *Segestria bavarica* and *S. florentina* seem to be closer to each other than to *S. senoculata*, but barcode data for *S. florentina* are not yet publicly available, and barcode distances indicate no especially close relationship between it and *S. bavarica*. The similarities between the two species could all be symplesiomorphic, but uniting them in the tree still seems the most plausible scenario.

### Tetragnathidae

The deep phylogeny of this family is based on the results of Kallal & Hormiga (2018). The relationships within *Metellina* and *Pachygnatha* are based on morphology and are strongly supported by analysis of the barcode sequences for all representatives. The same is the case for *Tetragnatha*, where the basal placement of *T. striata* is based on its previous position in a separate genus (*Arundognatha*) and its presumed relationship to *T. nitens* or *T. vermiformis* (Levi 1981). Barcodes for both of these have been sequenced and show a similar basal position relative to the remaining British congeneric species.

### Theridiidae

The phylogenetic framework for this family is based on the results of Liu et al. (2016), combining morphological and molecular evidence, with additional genera and species added according to the morphological analyses of Agnarsson (2004).

The genus *Steatoda* in the usual sense is clearly highly polyphyletic in the analyses by Liu et al. (2016); creating a monophyletic *Steatoda s. lat*. would require merging the genus with both *Crustulina* and *Latrodectus* (and possibly other Asagenini); certainly not a desirable solution. Instead, *Steatoda* is here tentatively divided into a number of smaller genera, all of which had been postulated previously, as classic authors have long recognised the heterogeneity of the genus. It is, however, important to realise that the molecular subdivisions are not closely aligned to previous morphology-based ideas (e.g., Wiehle 1937; Wunderlich 2008), and the two sets of results are difficult to reconcile.

Relationships within *Enoplognatha* are based on barcode data, *E. oelandica* being placed based on morphology (but with low confidence). The semispecies relationship between *E. ovata* and *E. latimana* is based on Lasut et al. (2015) who show convincing evidence based on mitochondrial and nuclear markers indicating that the two forms are not yet fully reproductively isolated, despite their obvious genitalic differences. Levi (1973) cites a personal communication from V. Seligy stating that both forms of the genitalia can be found among siblings from the same egg sac.

The internal topology of *Robertus* is also based on barcodes, but with weak support, the placement of *R. insignis* being based on its similarity to the barcode-sequenced *R. lyrifer* (Almquist 1978).

*Phycosoma inornatum* is here considered a member of *Lasaeola*, following Wunderlich (2020). This placement is problematic, given that both of the genera, as well as *Dipoena* seem to be poorly delimited. The transfer to *Lasaeola* is justified by the observation that European *“Phycosoma”* is unlikely to be congeneric with the type species from New Zealand: it lacks an epigynal scapus [present in true *Phycosoma*] and has a relatively large embolus and conductor [small in true *Phycosoma*]; the male prosoma is also very different and matches that of other *Lasaeola* species. The species thus lacks the most important genus diagnostic characters of *Phycosoma*. Otherwise, the arrangement of the species is maximally conservative and maintains *Dipoena* and *Lasaeola* as separate genera, despite concerns about their possible para- or polyphyly. Arrangements within each genus are based on morphological similarities (e.g., Miller 1967).

*Dipoena* (*Lasaeola*) *lugens* is a phantom species in the sense of Breitling et al. (2015, 2016), probably not native and not reported again since the original description. It is thus considered a *nomen dubium* and not included in the tree.

The relationships between the three *Episinus* species are inferred from their barcode sequences.

The placement of *Coleosoma floridanum* is uncertain, as it is not necessarily congeneric with the species analysed by Liu et al. (2016). The placement of *Simitidion* as sister of *Phylloneta* is based on Wunderlich (2008; general morphological affinity) and Knoflach (1996; mating behaviour). The placement of *Theonoe* is based on Agnarsson et al. (2007).

*Cryptachaea riparia* is considered a member of *Parasteatoda*, based on the arguments detailed in Breitling (2019d). The internal relationships between the *Parasteatoda* species are based on barcode data and morphology.

The relationships between the *Rugathodes* species are based on morphological similarities.

The genus *Theridion* is clearly polyphyletic on a global scale, and its phylogeny inferred on the basis of morphology and barcode data is not always consistent with more comprehensive molecular phylogenies. To retain monophyletic genera in the tree, *T. pinastri* is placed in *Allotheridion*, following Archer (1950): barcode data place the species very close to the type species, and the general genus concept proposed by Archer seems validated by the molecular data, including the closeness to *Phylloneta*. The transfer of *T. hannoniae* to the same genus is based on its membership in the *petraeum*-group (Bosmans et al. 1994). *Platnickina tincta* is returned to *Theridion s. str*., to be conservative and avoid changing the name of common species that have always been placed in *Theridion*.

### Thomisidae

Phylogenetic studies of the crab spiders, e.g., by Benjamin (2011), Benjamin et al. (2008), Ileperuma Arachchi & Benjamin (2019), and Ono (1988), focus on the higher-level phylogeny of the family and include relatively few species that have close relatives in the British fauna. Barcode data can help to guide tree reconstructions in areas left unresolved by these global analyses. *Misumena* is placed as sister to *Pistius*, based on the barcode data, with strong support. *Diaea* is considered sister of the two, based on morphological similarity, with *Thomisus* sister of all three. Arrangements within Coriarachnini are based on Breitling (2019a), placing *Bassaniodes* as sister of *Psammitis* + *Xysticus*, in a conservative arrangement relative to *Ozyptila* + *Cozyptila*. *Ozyptila sanctuaria* and *O. pullata* are considered sisters of *O. claveata*, based on their similarity to the barcode-sequenced *O. arctica*. Resolving the trichotomy at the basis of *Xysticus s. str*. in Breitling (2019a) is difficult with the available data, and *X. bifasciatus* is placed basal to the other British members of *Xysticus s.str*. without strong arguments in favour of this placement.

*Ozyptila maculosa* is a phantom species as defined by Breitling et al. (2015, 2016), possibly referring to a malformed specimen of *O. atomaria*. It is thus considered a *nomen dubium* and not included in the tree.

### Zodariidae

The proposed topology is based on the informal, non-cladistic species groups proposed by Bosmans (1997), which seem not entirely inconsistent with the limited barcode data available for the family.

## Discussion

The phylogenetic tree proposed here can be used for a large variety of applications, for instance in evolutionary and conservation biology. A few illustrative examples are shown in **Figures 2** and **3**. **Figure 2** illustrates the phylogenetic distribution of a number of classical morphological traits traditionally used in the identification of linyphiid spiders. The phylogenetic tree allows organising a large and unwieldy data matrix to facilitate the identification of patterns. While there is a high degree of homoplasy for all characters, many show clear trends in agreement with traditional (typological) classifications of Linyphiidae, such as the loss of tibial spines (Wiehle formula) and the distal movement of the trichobothrium on metatarsus 4 in “higher” linyphiids. In many cases, these trends extend beyond genus boundaries, and their identification requires the more detailed framework provided by the phylogenetic tree, e.g., regarding the the large body size in the “basal” Linyphiinae (*Linyphia, Frontinella* and their relatives). Another obvious use of this character map is its application as a starting point for correcting the tree and proposing better alternative hypotheses.

**Figure 3.**
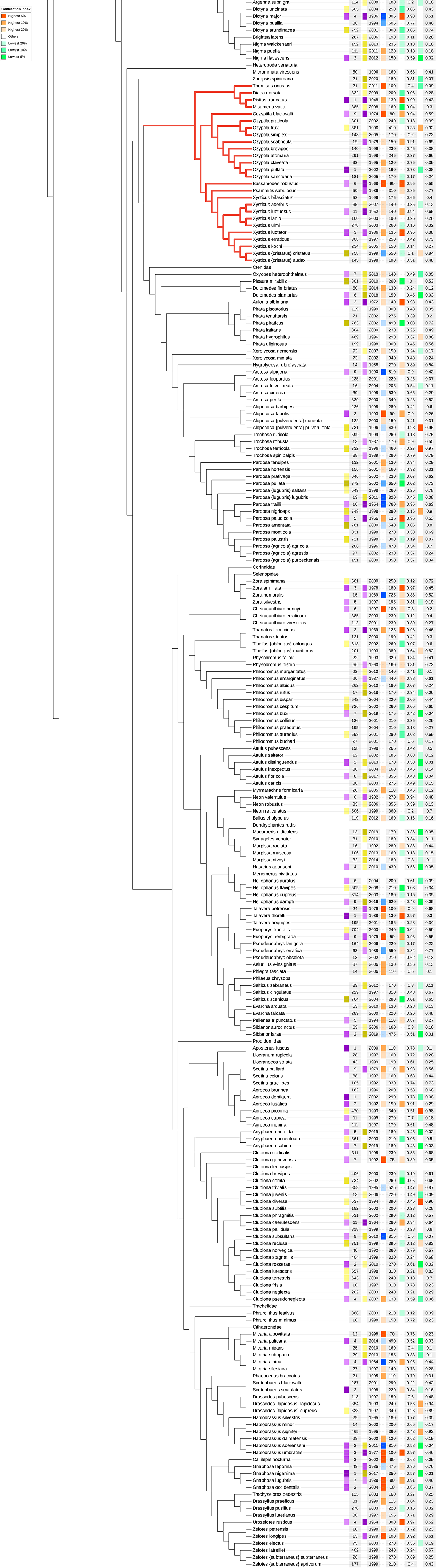
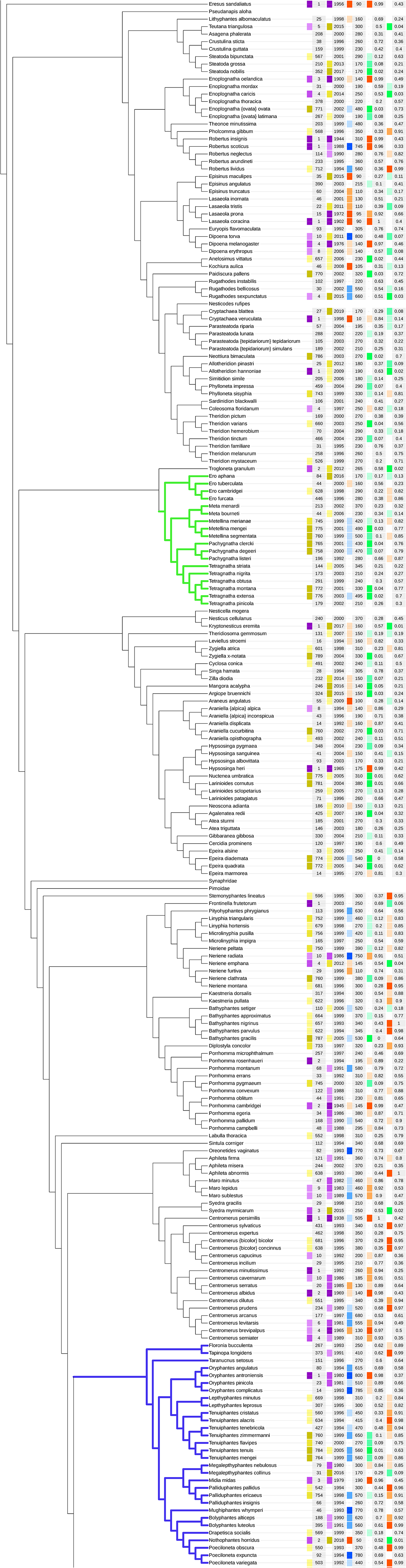
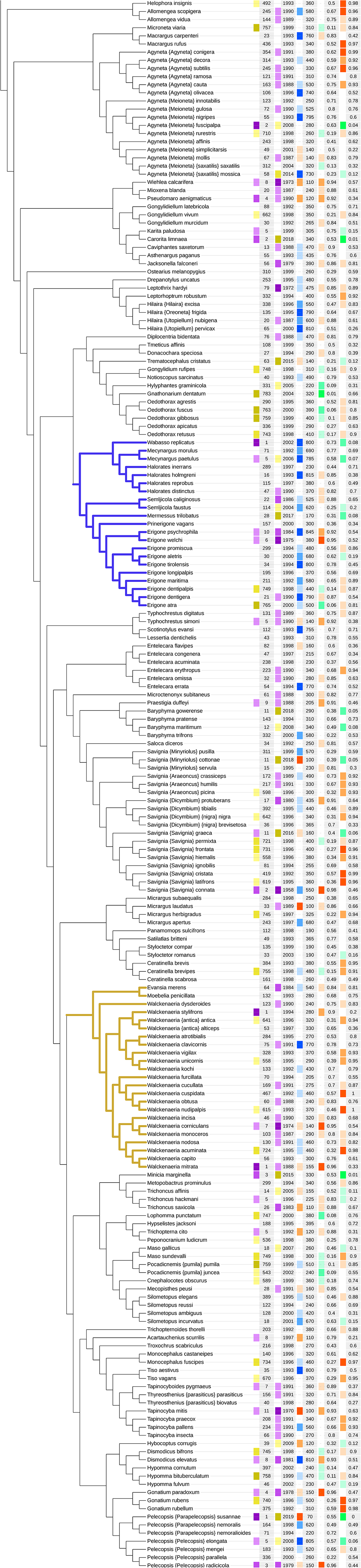
Selected ecological and conservation biological variables mapped onto the phylogenetic tree of British spiders. The abundance indicates the number of occupied hectads, with a 1% depreciation per year since the latest record. The recency of the observations is indicated as the median year of the latest record per hectad. The northern distribution is indicated by the median distance of the occupied hectads from the southern edge of the British Ordnance Survey national grid. The Vulnerability Index is 1 minus the quantile of the average of the recency quantile and the abundance quantile for each species. Higher values indicate species that are reported from only a small number of hectads and have few or no recent records; low values indicate species that are widespread and have often been recorded recently. The Contraction Index is the quantile of the difference between the abundance and recency quantiles. Higher values indicate widespread species with relatively few recent records; low values indicate highly localised species that have nevertheless a large fraction of recent records (e.g., new arrivals and expanding species). The clades highlighted in colour are discussed in more detail in the text. The order of columns (from left to right) follows the order of the legends (from top to bottom). Based on data provided by the UK Spider Recording Scheme.

**Figure 3** uses the tree to search for potential correlations between phylogeny and ecological traits. Data on the distribution of spiders in Great Britain (at the hectad level) were obtained from the Spider Recording Scheme database and a number of ecological indicator values were calculated for each species (excluding accidental introductions for which ecological analyses would be inappropriate): (1) an estimate of the abundance of the species, based on the number of occupied hectads; the weight of each hectad was decreased by 1% for each year since the latest record from this hectad; (2) an indication of the recency of observations, based on the median year of the latest record per hectad; (3) a characterisation of the North-South distribution based on the median distance of the occupied hectads from the southern edge of the British Ordnance Survey national grid; (4) a “Vulnerability Index”, 1 minus the quantile of the average of the recency quantile and the abundance quantile for each species, where high values (close to 1) indicate species that are reported from only a small number of hectads and have few or no recent records, while low values (close to 0) indicate species that are widespread and have often been recorded recently; (5) a “Contraction Index”, i.e., the quantile of the difference between the abundance and recency quantiles, where high values (close to 1) indicate widespread species, however with relatively few recent records, while low values (close to 0) indicate highly localised species that have nevertheless a large fraction of recent records (e.g., new arrivals and expanding species).

For each of the more than 680 subtrees a Wilcoxon rank sum test was applied to identify clades that are significantly enriched in members with particularly high or low values for each of these ecological indicators. In many cases, the results confirm previously reported informal observations. For instance, the clade most significantly enriched in species with a northerly distribution includes all the Linyphiidae compared to the rest of the species (Wilcoxon p-value < 5 × 10^−26^). However, the phylogenetic tree also allows an analysis at a much finer resolution.

In **Figure 3** a few selected examples are highlighted. Some of the results follow traditional family or genus boundaries and could be obtained without the help of a fully resolved tree. For example, the Thomisidae are significantly enriched in species with a southern distribution (Wilcoxon p-value < 5 × 10^−5^).

More interesting, however, are those trends that are only possible to identify within the frame-work of a fully resolved tree: for instance, the clade combining Mimetidae and Tetragnathidae is significantly enriched in species with a particularly low Vulnerability Index (Wilcoxon p-value < 5 × 10^−5^), i.e., these species are on average more widely distributed and more recently recorded than the members of other clades. The fully resolved tree also allows more detailed examination of trends within families, beyond genus boundaries. For example, species in the two clades from *Floronia bucculenta* to *Poeciloneta variegata* and from *Wabasso replicatus* to *Erigone atra* in **Figure 3** are highly enriched in species with a northern distribution (Wilcoxon p-values < 3 × 10^−5^ and < 3 × 10^-7^, respectively). This is one more example showing that ecological traits are correlated with phylogeny, beyond the coarse-grained categories of traditional Linnaean classification, as has been shown for other taxa before (e.g., Thomas 2008 for birds).

This is particularly interesting for those cases, where the ecological variables indicate potential conservation concerns. For instance, in **Figure 3**, the clade from *Evansia merens* to *Walckenaeria mitrata* is strongly enriched in species with a high Contraction Index (Wilcoxon p-value <5 × 10^−5^), i.e., these species show a surprising lack of recent records for such widespread species. There are numerous potential explanations for this phenomenon, not all of which indicate an immediate conservation concern, but this example nevertheless illustrates the potential of future applications of the fully resolved tree for spider conservation research.

In conclusion, it is perhaps useful to reflect on the degree of confidence in the presented tree. As stated repeatedly, there remain multiple areas where the available evidence does not yet allow a highly confident decision between alternative phylogenetic hypotheses. A fully resolved phylogenetic tree for the 680 British spider species could contain at most on the order of 650 “mistakes” relative to the true phylogenetic history of the group (the maximum number of branch moves required to transfer the tree into the correct one; Atkins & McDiarmid 2019). The expected number of mistakes in a random tree would be on the order of 600.

Of course, one would hope that the tree proposed here is far from random and provides a good initial approximation of the true phylogeny. Nevertheless, the number of mistakes is probably still considerable, given the remaining instability and incompleteness of the underlying datasets. This is a problem shared with almost all published phylogenies: for instance, Miller & Hormiga (2004) report that their analysis of erigonine phylogeny has only 5–6 nodes (about 20%) in common with the topology proposed for the same genera just 4 years earlier by Hormiga (2000), despite using very similar methodology. They suggest that 50–53 branch moves would be required to convert the trees into one another. Even if this appears to be an overestimate, considering the results of Atkins & McDiarmid (2019), it indicates that at least one of the trees was still very far from reconstructing the true evolutionary history of this small sample of the subfamily.

If the tree proposed here, for a much larger number of taxa, contains a similar number of mistakes, this could be considered a major success. The number of errors in the presented tree is difficult to estimate objectively, but the above calculations can provide a valuable point of reference: each reader is likely to find some parts of the tree where their personal interpretation of the data would lead them to prefer a different arrangement of the species. Each of them will be able to count how many corrections (“branch moves”) would be required to transform the presented tree into their preferred one. They can then compare the number of corrections to the number of errors expected in a random tree; the ratio between the two numbers provides a metric of the (subjective) correctness of the proposed phylogeny.

Of course, phylogenetic relationships for British spiders have been proposed before, not only for individual small groups, but implicitly also for the entire order, for instance in the arrangement of species in the works of, e.g., Locket & Millidge (1951, 1953) or Roberts (1985, 1987). But even when the proposals were made explicit in the form of tree diagrams, they sometimes failed to achieve the level of precision and specificity that would make the suggestions testable in an objective way. A good example is the tree proposed by Millidge at the conclusion of his analysis of the (mostly British) Linyphiidae (Millidge 1977:fig. 200): here, the tree not only consistently shows extant genus groups as “evolutionary precursors” of other groups and includes a number of tentative alternative branches, but most importantly it leaves a large number of unresolved polytomies. All this makes the proposals not only difficult to confirm or falsify, but challenges even the simple, direct comparison of individual proposals. This changes only when dichotomous trees are fully specified; this enabled, for instance, the detailed and quantitative comparison of alternative trees in Miller & Hormiga (2004). But even recent studies, which present fully resolved trees, often present multiple alternative topologies instead of identifying a single preferred tree, thus again reducing the resolution of the results by implying polytomies and diminishing the predictive content and testability of the hypotheses.

The explicit statement of all phylogenetic hypotheses in the form of a single completely resolved tree, i.e., a tree that only includes bifurcations and avoids polytomies, down to the level of individual species, makes the proposals of the present synthesis eminently testable. It is hoped that future analyses will identify errors and omissions in the presented tree and suggest alternative, better-supported hypotheses. At the same time, the tree might serve as a basis for extended analyses, perhaps initially extending the geographic scope to neighbouring countries, but ultimately resulting in a completely resolved tree of the global spider fauna.

